# The route of vaccine administration determines whether blood neutrophils undergo long-term phenotypic modifications

**DOI:** 10.1101/2021.10.26.465872

**Authors:** Yanis Feraoun, Jean-Louis Palgen, Candie Joly, Nicolas Tchitchek, Ernesto Marcos-Lopez, Nathalie Dereuddre-Bosquet, Anne-Sophie Gallouet, Vanessa Contreras, Yves Lévy, Frédéric Martinon, Roger Le Grand, Anne-Sophie Beignon

## Abstract

Innate immunity modulates adaptive immunity and defines the magnitude, quality, and longevity of antigen-specific T- and B- cell immune memory. Various vaccine and administration factors influence the immune response to vaccination, including the route of vaccine delivery.

We studied the dynamics of innate cell responses in blood using a preclinical model of non-human primates immunized with a live attenuated vaccinia virus, Modified vaccinia virus Ankara (MVA), and mass cytometry. We previously showed that MVA induces a strong, early, and transient innate response, but also late phenotypic modifications of blood myeloid cells after two months when injected subcutaneously. Here, we show that the early innate effector cell responses and plasma inflammatory cytokine profiles differ between subcutaneous and intradermal vaccine injection. Additionally, we show that the intradermal administration fails to induce more highly activated/mature neutrophils long after immunization, in contrast to subcutaneous administration.

Different batches of antibodies, staining protocols and generations of mass cytometers were used to generate the two datasets that were compared. Mass cytometry data were analyzed in parallel using the same analytical pipeline based on three successive clustering steps, including SPADE, and categorical heatmaps were compared using the Manhattan distance to measure the similarity between cell cluster phenotypes.

Overall, we show that the vaccine *per se* is not sufficient for the late phenotypic modifications of innate myeloid cells, which are evocative of innate immune training. Its route of administration is also crucial, likely by influencing the early innate response, and systemic inflammation, and the vaccine biodistribution.

## INTRODUCTION

Vaccination is among the major advances in terms of public health by conferring protection against many infectious diseases. However, there are still no vaccines against several pathogens, such as the human immunodeficiency virus (HIV). Furthermore, certain vaccines are still insufficiently effective, such as flu vaccines, which are not universal bu seasonal, or intradermal vaccination with *Mycobacterium bovis* bacillus Calmette-Guérin (BCG), which displays variable efficacy in preventing tuberculosis (TB) in adolescents and adults. These examples highlight the importance of carrying out in-depth investigations on the modes of action of vaccines to better understand the host and vaccine factors that influence innate and adaptive immune responses to vaccination (1) to guide and improve vaccine design.

The early innate immune response is not only among the first lines of antiviral defense but also orchestrates and modulates the antigen-specific effector and memory B- and T-cell responses by determining the frequency, functions, and dynamics of antigen-specific T and B cells (2,3). This is indeed the principle of vaccination.

The route of vaccine administration influences antigen/adjuvant trafficking, local and systemic inflammation, innate effectors, and the resulting adaptive immune response (4–7). We have shown that, like the antigen-specific antibody and T-cell responses, the early innate responses differ between subcutaneous (SC) and intradermal (ID) immunization, with the recruitment of distinct cell populations and activation of different immunomodulatory genes in skin and blood in response to a model live attenuated vaccine, Modified vaccinia virus Ankara (MVA), in non-human primates (NHPs) (8).

Here, we investigated the long-term impact of the route of vaccine delivery on the innate myeloid cell compartment. We hypothesized that the different early innate effector responses may lead to different innate immunological imprintings, which may last several weeks or months following vaccine injection. We previously showed that a SC injection of MVA induced late changes in the phenotype of innate myeloid cells in monkeys (9,10). They occurred between two weeks and two months after SC immunization, in spite of the resolution of systemic inflammation, shown by a return to baseline blood leukocyte counts and inflammatory cytokine levels. The innate myeloid response to a second MVA exposure two months later differed from the response to the first immunization, and to a second one two weeks later, in that it involved phenotypically more highly activated/mature neutrophils, monocytes, and dendritic cells (DCs). These late phenotypic modifications were associated with functional modifications as the *ex vivo* inflammatory response of PBMCs was enhanced short after the second immunization at two months compared to short after the first immunization and to a second immunization at two weeks (10).

We thus reused the mass cytometry dataset that originated from this previous preclinical study with macaques immunized with MVA SC (9,10) and compared the innate myeloid cell response with a newly generated dataset after ID injection. CYTOF data were analyzed similarly but independently, using sequential optimized clustering steps. They were compared using a strategy that we formerly developed that is based on categories of marker expression and Manhattan distance (10). Here, we demonstrate that ID administration of MVA failed to modify the phenotype of blood neutrophils long after immunization, contrary to SC injection, and although it mobilized neutrophils early after immunization.

## MATERIALS AND METHODS

### Animals

Adult male cynomolgus macaques (*Macaca fascicularis*) (n = 6, identified as 1BJR13, 1BJZ13, 1BLE13, 1GW14, AF103H and AN363H) were imported from Mauritius and housed in the CEA animal facility in Fontenay-aux-Roses, France.

### Vaccine, immunization and blood sampling

Animals were immunized twice, two months apart, with the ANRS recombinant MVA-HIV B vaccine (Transgene) at a dose of 4 × 10e^8^ plaque forming units (PFU) by ID injections distributed over 10 injection points (150 μL/point of injections, all along the back, in two columns). Recombinant HIV-1 antigens consisted of the complete sequence of gag, fused to fragments from pol (residues 172-219, 325-383 and 461-519) and nef (residues 66-147 and 182-206) of the Bru/Lai isolate. Blood samples were taken longitudinally before and after immunizations in either lithium-heparin (Greiner Bio-One) for mass cytometry analysis, or in ethylenediaminetetraacetic acid (EDTA) (Greiner Bio-One) for complete blood counts (CBCs) and plasma preparation. The same batch and dose of vaccine and vaccine schedule, but a different route of administration, were used relative to a previous preclinical study that analyzed innate responses after SC immunization (9), and for which the mass cytometry dataset was reused here.

### Determination of plasma antibody titers

Direct enzyme-linked immunosorbent assays (ELISA) were performed according to a previously published protocol (10) to determine total IgG titers specific to MVA in macaque EDTA-plasma. Antibody titers were calculated by extrapolation from a five-parameter logistic curve representing optical density (OD) versus plasma dilution and were defined as the reciprocal of the plasma dilution up to two times the OD of the plasma taken before corresponding immunization and diluted to 1:50.

Neutralizing antibody titers were evaluated using a previously described cell-based assay (10) based on the infection of HeLa cells with MVA-eGFP pre-incubated with serial dilutions of plasma-EDTA and flow cytometry analysis The curve representing the percentage of living eGFP^+^ cells as a function of the dilution of plasma-EDTA was plotted to calculate the neutralizing titer, equal to the reciprocal of the dilution of the sample resulting in two times less infected cells than after incubation with the plasma of the same macaque taken before immunization and diluted 1: 100.

### Complete blood count

Blood cell counts were determined from EDTA blood using an HMX A/L analyzer (Beckman Coulter).

### Measurement of plasma cytokine concentrations

The following cytokines, chemokines, and growth factors were quantified in plasma using a 23-plex MAP NHP immunoassay kit (PCYTMG-40K-PX30, Millipore) following the manufacturer’s recommendations: IL-17, GM-CSF, IFN-γ, IL1-β, IL1RA, IL-2, IL-4, IL-5, IL-6, IL-8, IL-10, IL-13, IL-12/23(p40), IL-15, IL-18, MCP1, MIP1-α, MIP1-b, scCD40L, TGF-α, TNF-α, VEGF, and G-CSF. Cytokine concentrations (in pg/mL) are plotted as a function of time (in days). The area under the curves (AUCs) of the early cytokine response after the first MVA immunization (H0 post-prime (PP) to D14PP) were calculated using GraphPad Prism 9 to represent the kinetics and magnitude of cytokine release as a single value. GM-CSF could not be quantified in plasma from SC-vaccinated animals. Heatmap was generated with DisplayR.

### Statistics

Ab titers, CBC and cytokine concentrations after ID immunizations were compared with Wilcoxon tests, whereas CBC after SC and ID injections were compared with unpaired two-tailed t tests, using GraphPad Prism 9.

### Leukocyte staining and acquisition by mass cytometry

A distinct, albeit highly similar, Ab panel (addition of 6 and suppression of 3 markers) and staining protocol (with or without heparin and with or without barcoding) were used between the ID and SC studies and the samples were acquired on a CyTOF 1 and Helios instrument for the SC and ID studies, respectively. Leukocytes were fixed extemporaneously without *ex vivo* restimulation and frozen following a previously described protocol that allows the analysis of all leukocytes and minimizes the batch effects inherent to the use of fresh cells (9). Briefly, three million fixed leukocytes were thawed and stained with a panel of Abs targeting innate myeloid cells (**Table S1**) in the presence of 300 U heparin to prevent nonspecific binding of metals by eosinophils (11). Purified Abs were conjugated to lanthanide metals using MAXPAR Lanthanide Staining kits (Fluidigm, South San Francisco, California, USA) following the recommendations of the supplier. Cells were barcoded using the Cell-ID 20-Plex Pd barcoding kit (Fluidigm). After washing in Barcode Perm Buffer, cells were incubated with one of the indicated combinations of Pd for 30 min at room temperature. Data were acquired using a Helios mass cytometer (Fluidigm) the day after staining after an overnight incubation in 0.1 μM iridium in PBS +1,6% PFA.

### Quality control and reproducibility

We controlled the quality and reproducibility of each staining and acquisition session (3 sessions in total with samples for all timepoints of interest from macaques 1BJR13 and 1BJZ13, 1BLE13 and 1GW14, and AF103H and AN363H) by staining and acquiring control samples consisting of aliquots of fixed and frozen blood leukocytes from a healthy macaque after *ex vivo* stimulation of whole blood for 2 hours with three TLR ligands: LPS (LPS *E. coli* 0111: B4, Invivogen) at 1 μg/mL, Poly I-C (Invivogen) at 100 μg/mL, and R848 (Mabtech) at 10 μM in the presence of Brefeldin A (10 μg/mL, SIGMA) for the last hour, in parallel with the tested samples. Comparison of the expression profiles of all markers by the control samples led us to exclude data from the first staining and acquisition session comprising the samples of two macaques (1BJR13 and 1BJZ13), as well as certain markers (CCR5, CXCR4, CD125, CD39, CD23, IL-1a) for which the expression profiles were too different from those of the two other staining and acquisitions sessions (**Figure S1**).

### Mass cytometry data preprocessing

Zero mean signal intensity (MSI) values were first randomized between −1 and 0 to avoid a bias in the density estimation by the SPADE algorithm. The FCS files were then normalized using the MATLAB program by Rachel Finck et al (12). Tube replicates were concatenated using the Cytobank tool (Mountain View, California, USA). Samples were de-barcoded using Debarcoder software (Fluidigm, San Francisco, USA) following the instructions of the user guide. The initial manual gating included the definition of singlets (based on Ir191/cell length), intact cells (Ir191/Ir193), no beads (Ce140/ Gd155), and the exclusion of CD3+CD66+ cells using Cytobank, as previously described (9). Although the use of heparin strongly reduced the nonspecific staining of eosinophils, a small number of CD3^+^CD66^+^ cells were still present and were excluded (approximately 0.2% of all acquired events).

### Automatic identification of cell populations with SPADE

The spanning-tree progression analysis of density-normalized events (SPADE) clustering algorithm (13) was used to automatically identify cell clusters or groups of cell clusters composed of phenotypically similar cells within the dataset consisting of singlet non-CD66^+^ CD3^+^ leukocytes from macaques (n = 4) collected at various timepoints (H0PP, H6PP, D14PP, H0 post-boost (PB), H6PB, and D14PB). Uniform random downsampling was used to select 100,000 cells from each sample (corresponding to the number of cells contained in the smallest sample) before a final upsampling step. The optimal SPADE parameters were determined using the SPADEVizR package (14) in R. The quality of the clustering was quantified as the percentage of clusters with a unimodal and narrow distribution for all markers of the classification. Marker distributions were assessed using the Hartigan’s dip test (p-value < 0.05 to reject the unimodality hypothesis). Markers distributions with an interquartile range < 2 were considered narrow.

### Heatmap representation of cell clusters

The SPADE tree was manually annotated based on the expression of key markers (CD1c, CD3, CD4, CD8, CD11c, CD14, CD16, CD20, CD66, CD123, CD125, CD172a, CADM1, and HLA-DR) to identify granulocyte clusters (**Figure S2**). A heatmap allowing easy visualization of the complete phenotype of each cluster of granulocytes was generated using the SPADEVizR by hierarchical clustering of clusters and markers based on the Euclidean metric (14). The range of expression of markers was divided between the 5^th^ and 95^th^ percentile into five categories for all clusters of cells. For each cluster, samples containing less than 50 cells were excluded from phenotype inference.

### Definition of phenotypic families

Clusters of cells sharing similar phenotypes were gathered into phenotypic families and superfamilies on the basis of the dendrogram resulting from the hierarchical clustering of the clusters, which were annotated manually. The number of cells for a given cell population was calculated as follows: CBC × population frequency.

### Phenotypic comparison of cell clusters

Both SC and ID datasets were analyzed independently using the same strategy. Clusters were compared two by two by calculating the Manhattan distance between the categories of expression of the clustering markers shared between the two datasets and visualized using the package CytoCompare in R (15).

## RESULTS

### MVA injected ID induces a strong specific humoral response

Adult macaques (n = 6) were immunized with 4 × 10e^8^ PFU of recMVA HIVB by ID injections twice, two months apart (**Figure 1A**). We assessed the humoral response to confirm the efficiency of vaccination, as Abs are the main immune correlate of protection for most vaccines (16), including vaccinia virus (VACV) and MVA against smallpox (17). As previously shown (8), MVA was highly immunogenic after ID injection. It induced high levels of MVA-specific IgGs (**Figure 1B**). As expected, the peak of the secondary response was higher than that of the primary response. Titers remained elevated, plateauing at 117,335 ± 50,060 28 days post-boost. Neutralizing activity was only detected after the second immunization (**Figure 1C**). The analysis was performed using EDTA-plasma, precluding a direct comparison with the Ab response induced in macaques identically immunized previously but through the SC route (10), which were analyzed using sera. However, in a previous comparative study, we reported that the adaptive responses differed between ID and SC MVA immunizations, with SC delivery inducing higher levels of nAbs, and ID delivery inducing more polyfunctional CD8^+^ T cells (8).

**Figure 1.**
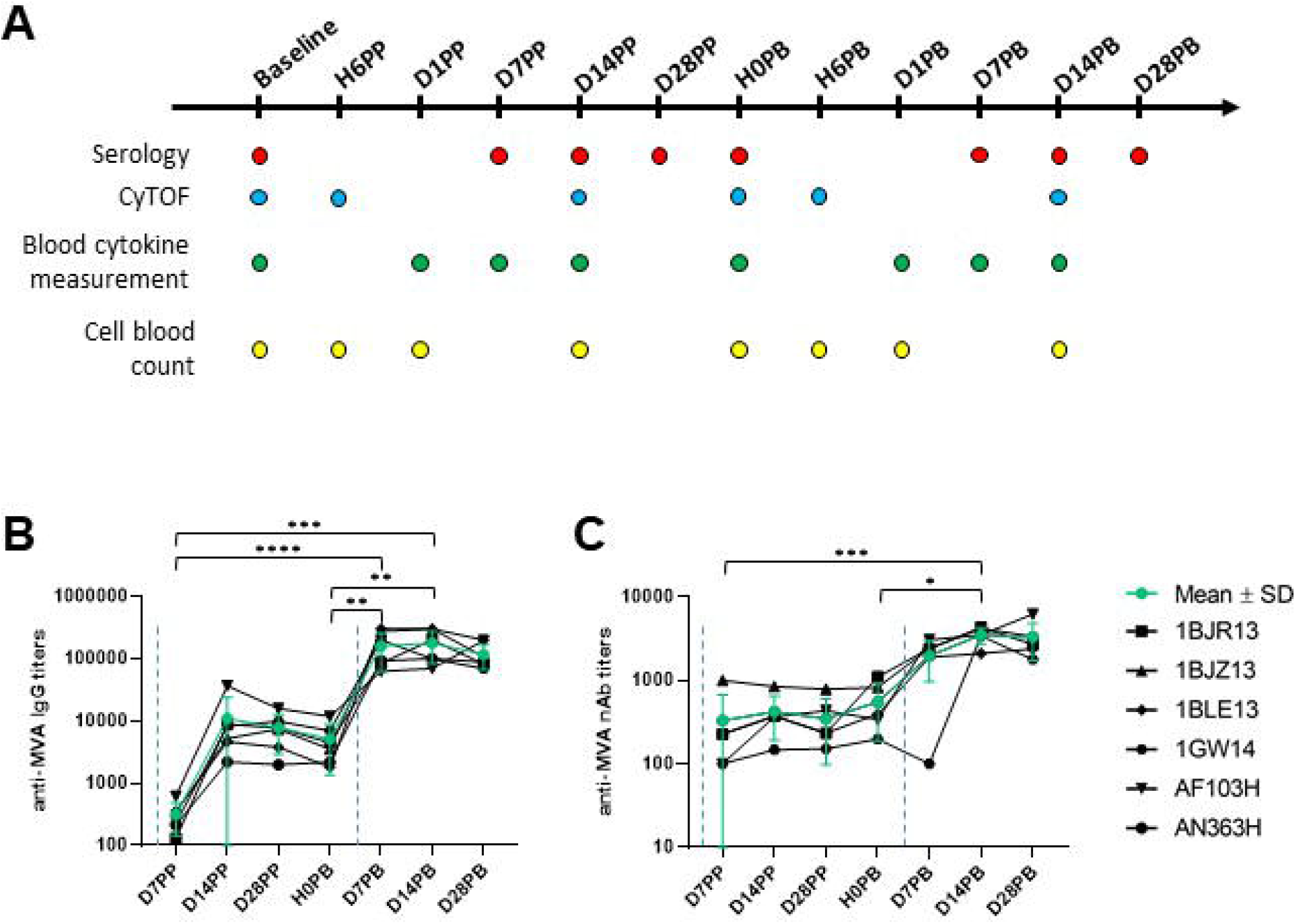
Experimental approach. **(A)** Experimental design. Six male and adult cynomolgus macaques were immunized two months apart with MVA HIV-B at a dose of 4 × 10e^8^ PFU injected intradermally. Blood was drawn longitudinally at hours (H for hour), days (D for day), or months (M for month) before and after the first (PP for post-prime) and second (PB for post-boost) immunization to assess the inflammatory, innate, and humoral responses and the phenotype of blood innate myeloid cells by mass cytometry. Immunizations are indicated by the blue dotted lines **(B)** Immunogenicity of MVA injected ID. Individual (black) and mean (green, with standard deviation) titers of anti-MVA IgG (left) and nAb (right) in EDTA plasma were plotted over time. Anti-MVA IgG titers were measured by ELISA. The MVA neutralizing capacity was quantified using a cell-based assay and MVA-eGFP. Immunizations are indicated by the blue dotted lines. Titers were compared by Wilcoxon tests. Statistically significant p values (p < 0.05) are indicated by an asterisk (*).

We next analyzed the innate immune response in blood overtime, as it shapes and can even predict the magnitude, quality, and persistence of the Ab response, with myeloid cells playing a key role in capturing and presenting vaccine antigens to B and T lymphocytes, we next analyzed them in blood over time.

### ID and SC immunizations with MVA induces strong but distinct inflammatory responses

ID injections induced a rapid and massive increase in the number of total leukocytes in blood (1.88 ± 0.29 times more leukocytes per milliliter of blood within 6 hours, p < 0.0001), but it was transient (the CBC decreased as soon as one day after immunization and returned to baseline 14 days post-immunization) and largely due to an increase in the number of granulocytes and, to a lesser extent, monocytes (**Figure 2A**). There was no difference between the first and second ID immunization (p = 0.95, paired t-test comparing the abundances at H6PP and H6PB). SC immunizations triggered a greater statistically significant increase in the total number of leukocytes, granulocytes, and monocytes (**Figure 2B**).

**Figure 2.**
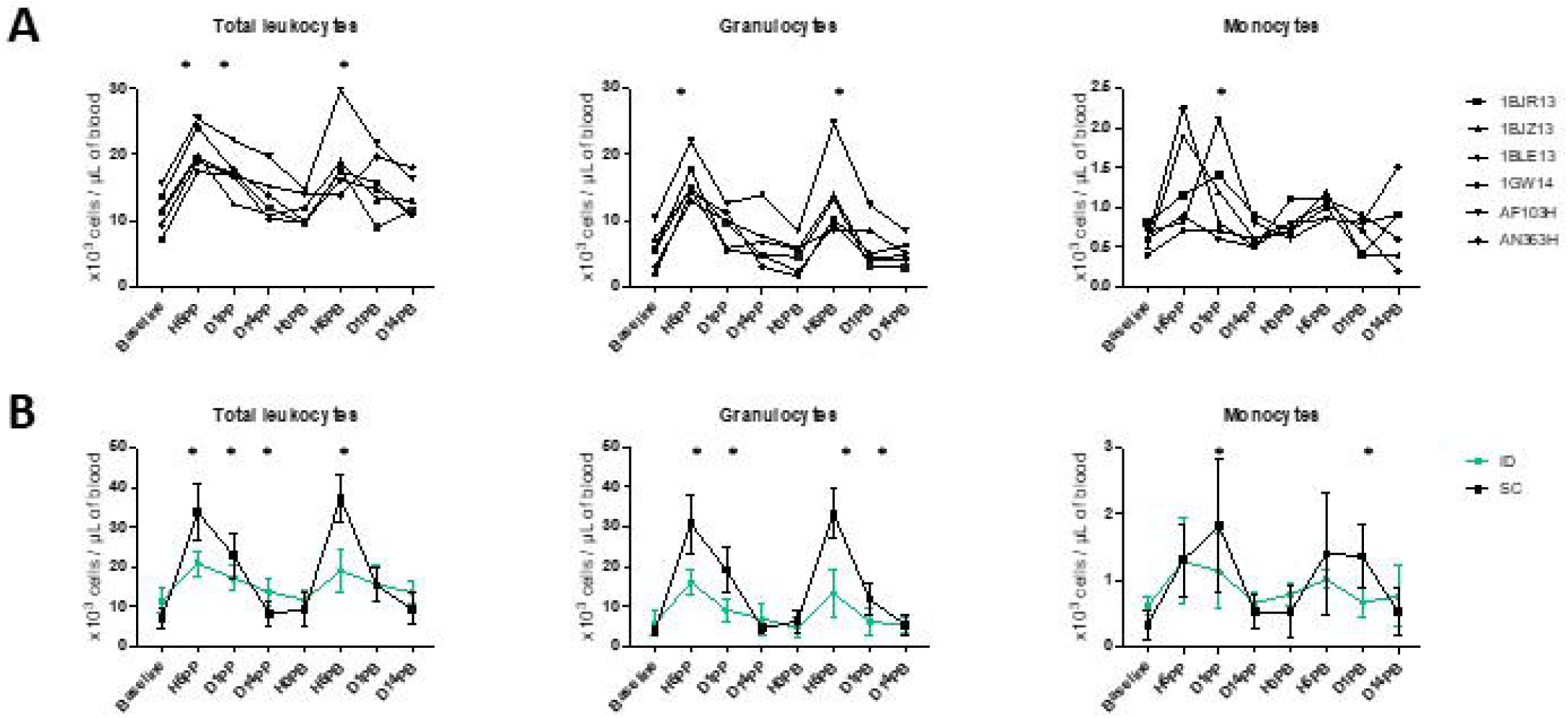
Leukocyte counts in blood after ID or SC immunizations. **(A)** Longitudinal monitoring of total leukocyte, granulocyte (including neutrophils, eosinophils and basophils), and monocyte counts (in thousands per μl of blood) before and after ID immunizations for each animal. Immunizations are indicated by the blue dotted lines. Counts were compared to baseline using Wilcoxon tests. p-values < 0.05 were considered statistically significant and are indicated by an asterisk (*). **(B)** Comparison of leukocyte counts (in thousands per μl of blood) before and after immunization by the ID (in green) or SC (in black) route. Immunizations are indicated by the blue dotted lines. Counts were compared between routes of vaccine delivery using unpaired two-tailed t tests. Statistically significant p values (p < 0.05) are indicated by an asterisk (*).

We next identified the systemic cytokine signature of MVA injected ID. The AUC, as an approximation of exposure over time, showed that, among 22 tested soluble factors, 16 cytokines were produced in response to the first ID injection: IL-12/23(p40), MCP-1, sCD40L, IL-1ra, VEGF, IL-10, IL-8, IL-13, G-CSF, IL-2, TNF-α, MIP-1α, IL-5, TGF-α, IL-4, and IL-17 (from the highest to the lowest cumulative concentrations) (**Figure 3A**). Only two cytokines differed significantly between the first and second ID immunization: MCP-1 concentrations decreased (p = 0.003) and those of IL-1b increased (p = 0.02). The cytokine profiles after the first ID or SC immunization differed quantitatively and qualitatively (**Figures 3B and 3C**). Immunization by the ID route induced a 1.79 times greater production of cytokines than that by the SC route (p = 0.009; Wilcoxon tests), with five cytokines representing more than 90% of those produced (**Figure 3B**), whereas immunization by the SC route resulted in a more diverse cytokine production profile, with five additional cytokines: MIP-1 β, IL-15, IFN-γ, IL-6, and IL-1 β (**Figures 3B**). Unsupervised hierarchical clustering showed SC-immunized animals to be highly similar to each another in terms of cytokine production, in contrast to ID-immunized animals, which showed greater heterogeneity(**Figure 3C**). Based on the cytokine dendrogram, the following cytokines distinguished between the two routes of MVA delivery: IL-15, IFN-γ, MIP-1 β, IL-6, TGF-α, IL-1 β, IL-4, IL-5, IL-2, TNF-α, MIP-1α, IL-17, and G-CSF (**Figure 3C**).

**Figure 3.**
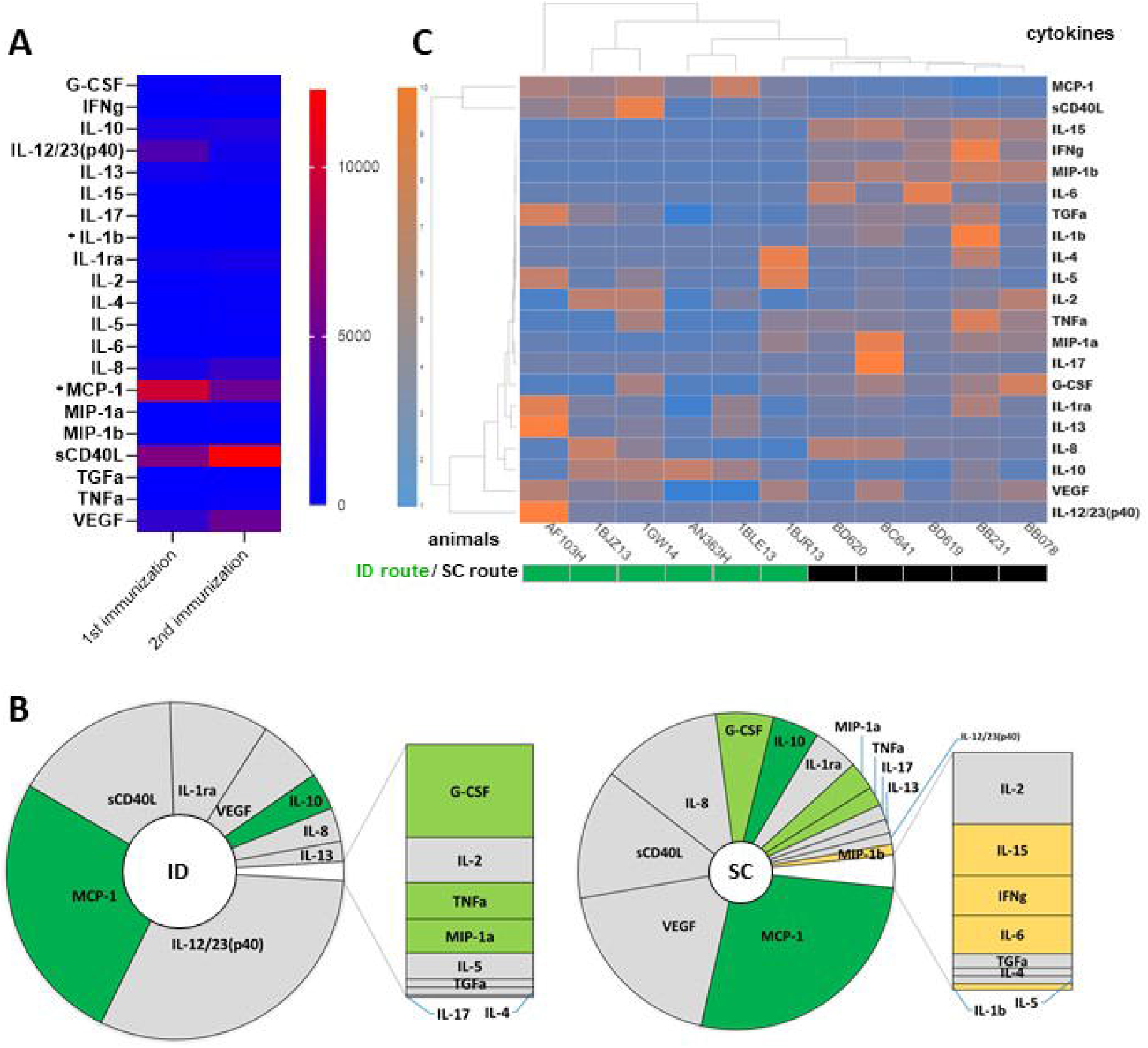
Comparison of cytokine and chemokine expression profiles in blood after ID or SC MVA immunizations. The plasma concentration of 22 cytokines was measured before, 1 day, 7 days, and 14 days after MVA injected either ID (n = 6) or SC (n = 5). Areas under the curve from H0 to D14 post-immunization were calculated to represent an approximation of exposure over time, in pg×ml^-1^×day. **(A)** The mean AUC of each cytokine after the first and second MVA ID injection is shown as a heatmap. Colors represent the AUC values from the lowest (blue) to the highest (red), standardized for each row. Statistically significant differences (p < 0.05, Wilcoxon tests) between the 1^st^ and 2^nd^ injection are indicated by an asterisk (*). **(B)** The mean AUC of each cytokine produced after the first ID (left) or SC (right) MVA injection is displayed as a pie-chart. Differentially expressed cytokines according to the vaccine delivery route are colored in dark green for cytokines with a higher concentration after ID injection than SC, in light green for those with a higher concentration after SC injection than ID, and in yellow for those produced only after SC injection. The size of the inner pie is proportional to the sum of the AUC of all cytokines. **(C)** Heatmap representation of individual AUC for each cytokine after the first ID or SC injection of MVA. Each row corresponds to a cytokine and each column to an animal. Dendrograms represent the hierarchal clustering of animals (upper) and cytokines (left) based on the Euclidean distance using the Ward2 clustering method. Colors represent the AUC values from the lowest (blue) to the highest (orange), standardized for each row.

Overall, our results showed that MVA injected ID led to strong systemic inflammation, which was resolved rapidly. The inflammatory response was comparable between the first and second ID injection, except for IL-1b and MCP-1, but different from that following SC injection in terms of the magnitude of the cellular response and cytokine profile in blood. We next extensively characterized the phenotype of innate blood myeloid cells over time after MVA ID immunizations.

### Blood granulocytes are highly heterogeneous and diverse

Blood leukocytes were stained with iridium and a panel of 35 titrated lanthanide-conjugated antibodies targeting innate myeloid cells, including granulocytes (**Figure 4A and Table S1**), and analyzed by mass cytometry. We used a previously described analysis pipeline (9) comprised of several successive clustering steps to identify cell clusters and groups of cell clusters with a similar phenotype and characterize their dynamics in response to immunizations (**Figure 4B**). The SPADE algorithm was first applied to all samples of the dataset. It consisted of 26 samples, corresponding to four of the six immunized macaques (1GW14, AF163H, AN363H, 1BLE13) and six time points (H0PP, H6PP, D14PP, H0PB, H6PB, and D14PB) (**Figure 1A**), minus two samples that were not available for technical reasons (H0PP for macaque 1BLE13 and D14PP for macaque 1GW14).

**Figure 4.**
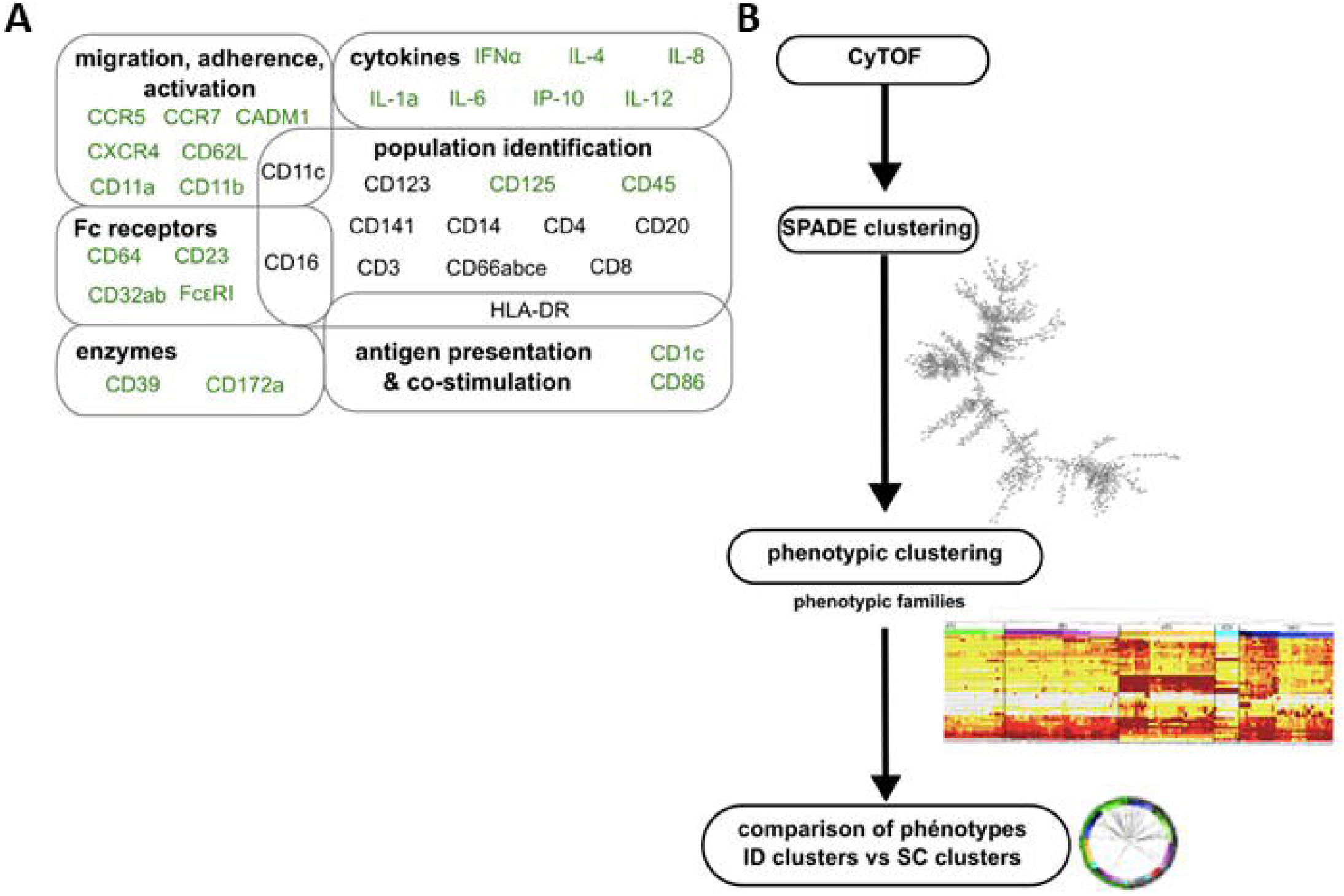
Analytical approach for mass cytometry characterization of granulocytes. **(A)** Panel of antibodies for mass cytometry analysis. Fixed leukocytes were stained with a panel of metal-conjugated antibodies (**Table S1**). Markers in black were used to compare the phenotype of clusters between datasets. Markers in light grey were excluded from the analysis, not selected as SPADE clustering markers for the ID dataset, or not shared with the SC dataset. **(B)** Analysis pipeline of mass cytometry data. After ID immunizations, cells sharing a similar phenotype were clustered using the SPADE algorithm. Clusters were manually annotated on the SPADE tree, and granulocytes clusters identified (**Figure S2**), and further grouped into “phenotypic families” and “superfamilies” after categorization of their marker expression and hierarchical clustering. Finally, a comparison of marker expression categories based on the Manhattan distance was used to associate granulocytes from the two datasets analyzed separately that display a similar phenotype.

We compared various parameters of SPADE, with the following parameters found to be optimal: 800 clusters, 22 clustering markers (CD66, HLA-DR, CD3, CD64, CD8, CD123, CD11a, CD11b, CD62L, CD4, FcεRI, CD86, CD172a, CD1c, CD32, CD16, CD11c, CD14, CD141, CD20, CCR7, and CADM1), and a downsampling of 20%, enabling maximum quality clustering, with 58% of the clusters having a uniform and narrow distribution of all clustering markers. As expected, these parameters differed from those of the SC dataset (9). Cell clusters on the resulting SPADE tree cell were manually annotated based on the expression of several key markers (CD66, CD3, HLA-DR, CD8, CD123, CD4, CD125, CD172a, CD1c, CD16, CD11c, CD14, CD141, CD20, and CADM1) and the major leukocyte populations (B cells, T cells, NK cells, monocytes and DCs, and granulocytes) were identified (**Figure S2**).

We next focused the analysis on the granulocyte compartment, as our panel was solely dedicated to target innate myeloid cells, and because major phenotypic modifications occur mainly in granulocytes late after MVA SC immunizations (18). The phenotypes of the granulocytes involved in the vaccine response were organized in the form of a heatmap after hierarchical clustering of the cell clusters and markers, once the levels of expression were categorized into five classes of signal intensity (**Figure 5A**), to visualize them more easily. Based on the clusters dendrogram, clusters sharing a similar phenotype were grouped into so-called “phenotypic families” and further grouped into “superfamilies”. Sixteen distinct phenotypic families (1 to 16) were distinguishable and grouped into five superfamilies (A to E). Neutrophils (CD66^hi^) were clustered within three superfamilies (A, B and E) according to their relative level of activation within the dataset. Superfamily A (phenotypic families 1, 2, and 3) comprised the least activated neutrophils of the dataset, with very low to low expression of cytokines (especially IL-8) and activation markers (especially CD45). Superfamily B (phenotypic families 4, 5, and 6) was composed of moderately activated neutrophils, with stronger expression of IL-8, CD86, CD39, CADM1, CD45, and CD11b, compared to superfamily A. Of note, among superfamily B, family 5 showed high expression of CD14, a characteristic already reported for neutrophils of cynomolgus macaques, without its function being clearly defined (19). Superfamily E (families 11, 12, 13, 14, and 15) contained the most highly activated neutrophils of this dataset, which expressed activation and migration markers and cytokines. This superfamily showed strong heterogeneity, from neutrophils of families 11 and 12 with the highest expression of activation markers, including IL-8, CD14, CD11a, CD11c, CD141, CD16, CD86, and HLA-DR, to neutrophils from family 13 displaying a relatively less activated or mature profile.

**Figure 5.**
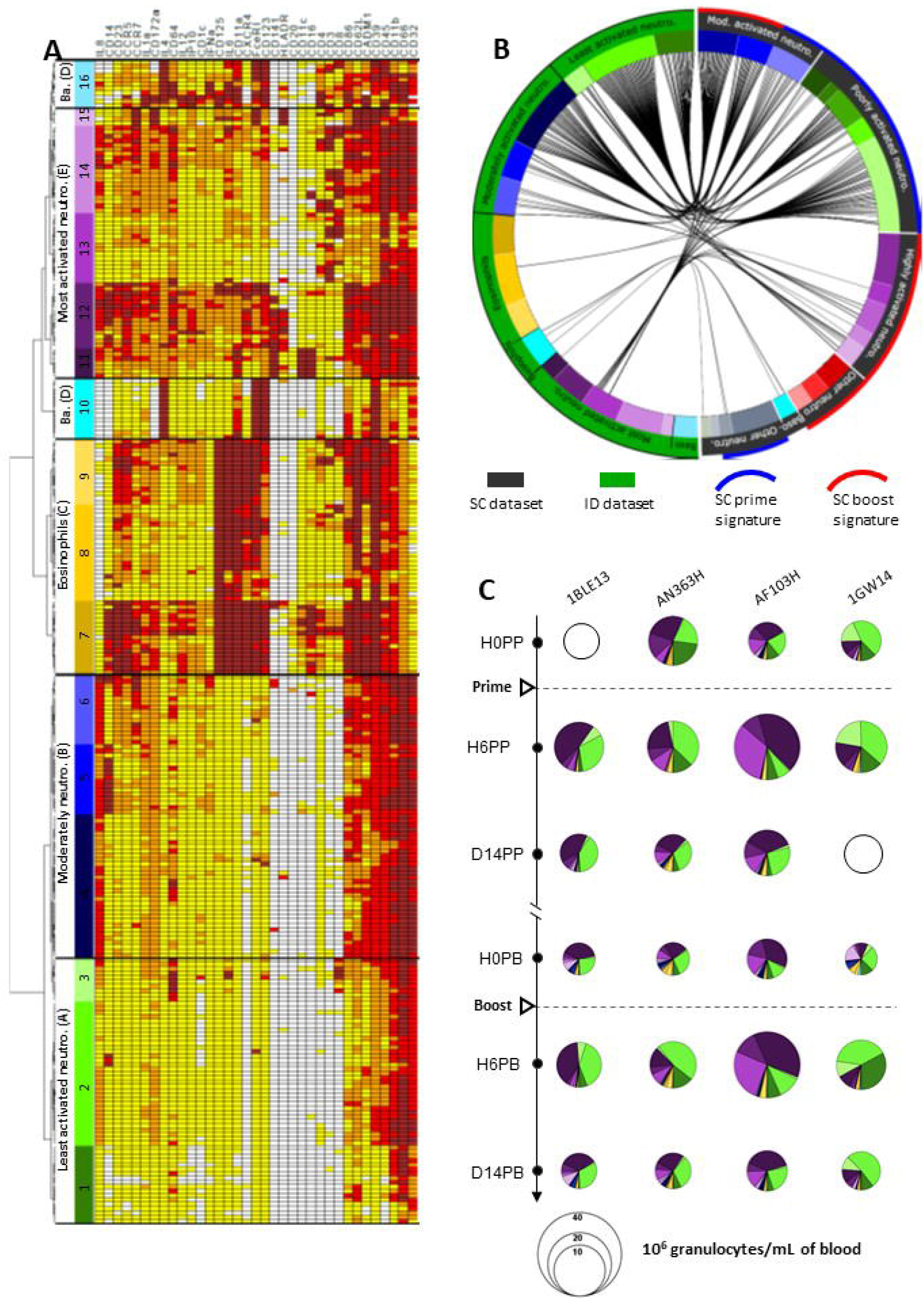
Phenotypic diversity and dynamics in granulocytes after ID and SC MVA immunizations. **(A)** Hierarchical clustering of granulocyte clusters represented as a heatmap. Each row corresponds to a cell cluster and each column to a marker. The level of expression of the markers was divided into five categories ranging from white to brown. The dendrogram allowed the grouping of clusters with similar phenotypes into 16 “phenotypic families” (numbered from 1 to 16) and “superfamilies” (named with letters between brakets, from A to E), colored based on the manual annotation as in (9). Neutro.: neutrophils. Ba.: basophils. **(B)** Comparisons of the phenotypes of granulocytes present in the ID (green) and SC (black) datasets. Black lines inside the circle connect phenotypically similar clusters after calculation of the Manhattan distance. The color code is identical to that of the heatmap. Neutro.: neutrophil. Mod.: moderately. Baso: basophils. **(C)** Pie charts representing the composition in granulocyte phenotypic families over time for each macaque after MVA immunizations by the ID route. Each slice represents a phenotypic family, for which the color is identical to that of the heatmap. The size of the pie is proportional to the cell concentration in the blood. Unavailable data are represented by empty circles.

Superfamily C (phenotypic families 7, 8, and 9) was clearly separated from the rest of the granulocytes. It included eosinophils with a CD66^med^ CD125^+^ phenotype and high expression of several markers, including CD45, CD62L, CD11a, CD125, CD23, IL-6, CXCR4, and FceRI, suggesting strong activation, especially for family 7. Nevertheless, these results must be interpreted with caution. The use of heparin during staining may have insufficiently inhibited interactions between the content of the eosinophilic granules and the metals conjugated to the Abs, and thus may have insufficiently prevented artefactual positive staining (11), making the phenotypic characterization of eosinophils by mass cytometry difficult.

Finally, basophils (CD123^+^ HLA-DR^-^) were found in superfamily D (phenotypic families 10 and 16). Family 10 gathered basophils with a classic expression profile (CD123^+^ IL-4^+^ FceRI^+^), whereas family 16 grouped basophils with more diverse expression profiles.

Thus, we show a large phenotypic heterogeneity and diversity of blood macaque granulocytes, as previously reported (9). We next investigated whether the phenotypic diversity within the ID dataset was qualitatively close to, or different from that of the SC dataset.

### The ID and SC datasets share similar poorly and moderately, but not highly, activated neutrophils

The analytical challenge for the phenotypic comparison between our two datasets lay in the use of different, albeit similar, Ab panels, Ab batches, staining protocols, and generations of mass cytometer, with different detection sensitivity. In addition, independent analyse were performed, following the same workflow, but using different SPADE parameters (cluster numbers, clustering markers, and downsampling), which were optimally defined for each dataset. We opted for a phenotypic comparison of the cell clusters based on the Manhattan distance between the expression categories of the clustering markers shared between the two datasets to address this challenge. The proximity between cell cluster phenotypes is shown in the form of a circular graph after calculating for the sum of the absolute value of the difference between categories for each shared marker for each paired clusters. This distance was penalized when one of the terms was greater than 1. Distances equal to or less than 11 were considered as significant, and are represented by linking the two compared cell clusters (**Figure 5B**).

As expected, the eosinophils (superfamily C) from the ID dataset could not be associated with any cluster from the SC dataset, as they were removed from the analysis due to of their nonspecific staining in the absence of heparin during staining. Conversely, we found associations between basophils from the ID and SC datasets, in particular from family 10. More importantly, only the least activated and moderately activated neutrophils from the ID dataset were associated with neutrophils from the SC dataset, specifically with the poorly and moderately activated neutrophil families. On the contrary, the vast majority of the most highly activated neutrophils from the ID dataset were not associated with any neutrophil cluster of the SC dataset. Only neutrophils from family 13 from the ID dataset were associated with neutrophils of the SC dataset, either with moderately or highly activated neutrophils. As previously shown, these neutrophil clusters did not exist prior to SC immunization. They were induced long after MVA prime immunization (9) and responded to the second immunization. They were part of the SC boost signature defined in the LASSO-LDA model (9).

Such a phenotype-limited comparison provides information about the presence of shared (basophils and poorly and moderately activated neutrophils) or specific (highly activated neutrophils) cell populations between the two datasets, both showing a high diversity of phenotypes, but does not indicate when, or in what proportion these granulocytes circulate in blood before and after ID *versus* SC immunizations.

### ID injection does not result in late modifications of the blood neutrophil phenotype, in contrast to SC injection

We determined the impact of ID injections on granulocyte populations by investigating the differences in their cell abundance over time represented as pie-charts (**Figure 5C**). As classically shown, eosinophils and basophils were the least numerous granulocytes present at baseline and did not show major changes in frequency after the first or second immunization. Among neutrophils, clusters from two superfamilies, the least (families 1, 2, and 3, which matched families 5, 7, 3, 1, 11, 13, and 4 in the SC dataset, **Figure 5B**) and most highly activated neutrophils (families 11 and 12 which were not phenotypically associated with neutrophils within the SC dataset, **Figure 5B**) were found at baseline. They represented the majority of cell types. Thus, the steady-state differed between studies, as neutrophils present before SC immunization were poorly and moderately activated (9). Six hours after the first ID immunization, the number of granulocytes increased, without major modifications in the proportion of the various subpopulations, still with a predominance of the least and most highly activated neutrophils. On day 14, cell counts returned to baseline values. Hence, the early response of granulocytes differed between ID and SC in magnitude and composition, but from the beginning. Long after the first ID immunization, and immediately before the second, the granulocyte composition showed slight changes, whereas the cell counts remained at the basal level. The frequency of the least activated neutrophils was slightly lower, in favor of the most highly activated neutrophils and, to a lesser extent, moderately activated neutrophils, which were almost absent at baseline, although the difference was not statistically significant. This redistribution was not commensurate with the major modifications of composition seen long after SC immunization, with the appearance of highly activated neutrophils (9,10), which had no counterpart in the ID dataset (**Figure 5B**). Finally, the neutrophil responses to the first and second ID immunizations highly resembled each other, in sharp contrast to what was observed after SC immunizations (9,10).

## DISCUSSION

### ID administration of MVA fails to induce neutrophils that are more highly activated/mature long after immunization and that respond to a second immunization, in contrast to SC

In-depth phenotyping by mass cytometry combined with an analysis pipeline composed of successive clustering steps, specifically and previously developed for longitudinal multidimensional data, allowed us to analyze the quantitative and qualitative differences between the innate myeloid cell responses in blood after one or two ID immunizations of MVA and to compare them to those following another route of immunization, SC, using the Manhattan distance to measure the similarities of phenotype between cell clusters. MVA administration induced the substantial, rapid, and transient recruitment of granulocytes to blood, regardless of its route of delivery. However, such early mobilization was stronger after SC injection of the vaccine. Granulocyte counts increased, whereas the cell subset composition remained unchanged early after the first ID and SC immunization, mirroring that present before immunization. The early response to a second ID injection of the vaccine did not differ from the first in terms of magnitude and dynamics. Most importantly, it involved a similar distribution of cell subsets, in contrast to the response to a second SC injection, which engaged more highly mature/activated cells that were induced long after the first administration of the vaccine (9). Thus, depending on the route of MVA administration, not only the Ab and T cells responses (8), and the early inflammatory/innate responses to the first immunization (**Figures 2B, 3 and 5C**, and as previously shown (8,9)) differed, but also the long-term impact on neutrophils (**Figure 5C** and as previously shown (9)).

### Comparing high dimensional cytometry datasets

High-dimensional cytometry, including mass cytometry and spectral flow cytometry, is a powerful technique to monitor immunity, identify cell subsets associated with diseases or potent responses to vaccines, and decipher the complex mechanisms of the immune response to vaccines (20–27). However, the comparison and integration of different datasets generated at multiple sites and on different days is challenging, even when using the same technology. This is due to technical differences, called batch effects, that affect the signal intensity (on which commonly used unsupervised analytical methods, such as SPADE, visNE, FlowSOM, CITRUS, and UMAP, are based) and need to be distinguished from true biological variability (28). Several algorithms have been proposed to normalize signal intensity to reduce batch effects before unsupervised cell cluster identification and to compare multiple datasets, such as CytofRUV (29), and JSOM (30). iMUBAC can even compare different datasets in the absence of shared technical replicates, used as reference samples, by overlaying cells from several healthy controls as anchors (31).

The issue here is somewhat distinct, as the two datasets were analyzed separately. The raw mass cytometry data generated after the SC immunizations were not re-analyzed together with the mass cytometry data newly generated after the ID immunizations, and after batch effect correction. High intra-cluster homogeneity was achieved using the SPADEVizR package for each independent analysis. Then, cells from the two vaccine immunogenicity studies (SC vs ID vaccine administration) were matched by comparing their categories of marker expression, instead of the mean intensity, to mitigate the expected technical differences in staining efficacy and cytometer sensitivity. Five categories were defined based on the range of expression of each marker between the 5^th^ and 95^th^ percentiles for each dataset. Distances were calculated as the sum of the absolute value of the Manhattan distance between the categorical values for each shared marker of each cluster. Two clusters were associated if the sum was below a certain threshold, and a penalty was applied, if the value of a term was too high. The threshold and penalty were set after trial and error and manual inspection of the heatmap and marker expression profiles between clusters. Given the large differences in the neutrophil response to MVA between the two delivery routes, such a simple method, and admittedly not scalable nor benchmarked against other algorithms, such as overlaying cell clusters onto a reference scaffold map (32), or measuring the quadratic form distance post-clustering (33), proved to be sufficient to define a stringent cluster-wise comparison.

### Neutrophils and the humoral response to vaccines

Neutrophils are the most numerous leukocytes in the blood. Their unexpected phenotypic and functional heterogeneity, and plasticity have been recently reported (34). Apart from their key antimicrobial activity, they shape adaptive immunity (35,36) by releasing cytokines and chemokines, granules, and NETs and by interacting with other immune cells and eventually acting as antigen presenting cells (APC) (37,38). Neutrophils can directly provide B-cell help through the production of BAFF and, hence, contribute to plasma-cell generation and antigen-specific Ab production (39–41). Whether and how neutrophil subsets, including highly activated neutrophils induced long after MVA SC injection, interact with B cells and modulate the primary and secondary MVA-specific Ab response is yet to be determined.

### Innate immune training and vaccines

Certain vaccines, in particular certain live attenuated vaccines, such as BCG and oral poliomyelitis vaccine (OPV), also provide protection against unrelated pathogens beyond the pathogen they initially target by a process called nonspecific effects of vaccines (NSE) (42). Cross-reactive T and B cells and antibodies (Abs), bystander activation, and innate immune memory, also called trained immunity, contribute to NSE. By definition, trained cells respond faster and more strongly to a secondary challenge with homologous or even heterologous pathogens than a ‘naïve’ cells, long after the initial stimulation (43). The mechanisms of trained immunity involve the epigenetic reprogramming of innate cells, which are known to be short-lived, and that of their hematopoietic stem and progenitor cells (HSPCs) in the bone marrow (BM). Resting trained cells display an enhanced responsiveness to a challenge without having themselves encountered the trained immunity-inducing stimulus, by inheritance from their HSPCs. BCG is a canonical trained immunity stimulus. It induces trained monocytes and neutrophils (44–46) that contribute to its anti-TB and nonspecific protective effects (47). In addition to BCG, several microbial, as well as endogenous, ligands have been shown to imprint innate cells, with increased pro- or anti-inflammatory responsiveness that may be beneficial or detrimental to the host (18).

SC injection of MVA, but not ID, induced the late presence of highly activated neutrophils that were better equipped to respond to a second injection. This raises the questions of whether these cells are authentic trained cells, how they compare with BCG-induced innate memory cells in terms of features and origin, and how the route of MVA delivery influence their generation.

The literature has emphasized the enhanced responsiveness of blood trained cells to challenge through the persistence of epigenetic marks inherited from trained HSPCs. However, the modulation of expression of certain markers associated with increased activation by resting trained but unchallenged cells, including neutrophils, has been reported after BCG given ID to humans (46,48). The phenotypic modifications of neutrophils induced by MVA injected SC are yet to be associated with functional and epigenetic modifications.

The state of neutrophil activation prior to immunization differed between animals from the two cohorts, which were otherwise healthy, without any low-grade inflammation and with CBCs in the normal range. This is likely the result of their different immunological history related to different environmental exposure, although they were naive (*i*.*e*., not previously experimentally immunized nor infected) (49). As recently stated, to the degree that vaccine-induced trained immunity can provide heterologous protection, the trained immune status at the time of vaccination may also modulate the immunogenicity of vaccines (50), including their capacity to further train innate cells.

### Route of vaccine delivery and innate immune responses

The route of injection determines the biodistribution of vaccines and their kinetics of expression, including those of VACV and MVA. In mice, MVA-expressing neutrophils were shown to migrate from the skin to the BM following ID injection (54). VACV can infect human BM hematopoietic stem cells *in vitro* (55). Intraperitoneal VACV injection in mice was shown to induce the rapid and transient expansion of HSPCs, with a bias towards common myeloid progenitors (CMP), which was MyD88 dependent. Finally, the route of MVA delivery modulates the early systemic inflammatory cytokine response, as reported here (**Figures 3B and 3C**). Not only did the profile of cytokines produced early in response to ID or SC delivery of MVA differ, but interestingly, IL-1 β and IFN-γ, proposed to play a key role in the induction of trained immunity (44,46,57,58), were differentially induced depending on the route of MVA delivery. Both were produced at low levels relative to those of other cytokines, and exclusively after SC injection. Whether MVA needs to reach and reside in the BM for a while, to be directly detected by HSPCs, or even infect them, to induce the late modifications of neutrophil phenotype is not known.

BCG has previously been reported to induce trained monocytes in mice after intravenous (IV) injection,, but not after SC injection. BCG was present in the BM for up to seven months, where it infected monocytes/macrophages but not HSPCs after IV injection, whereas it was absent from the BM after SC injection. The long-term persistence of BCG in the BM was not required for the induction of immune training. The almost complete clearance of BCG from the BM by a four weeks antibiotic treatment with antimycobacterial drugs four weeks after IV injection did not prevent BM Lin-Sca1+c-Kit+ (LSK) cells to expand (44). In macaques, pulmonary mucosal delivery of BCG was also shown to induce trained monocytes in the blood and BM in more vaccinated animals than “classical” ID injection (50). However, whether the early systemic inflammatory/innate responses also contribute to the difference between IV and SC injection, or ID and mucosal delivery for innate immune training was not addressed.

Future studies are needed to fully understand which vaccines/adjuvants imprint innate cells long after their administration and how, in particular by defining the respective roles of the vaccine biodistribution and persistence, and the early inflammatory/innate response, to harness this property to optimize protection against infectious diseases, whether it is mediated by the innate or adaptive immune systems (18,59,60).

## DATA AVAILABILITY STATEMENT

Mass cytometry data were deposited publicly. FCS files from the SC and ID studies are available on the FlowRepository database through ID FR-FCM-ZYBG and FR-FCM-Z4KJ, respectively.

## ETHICS STATEMENT

The study was approved by the Ministry of National Education, Higher Education and Research and by the Ethics Committee in Animal Experimentation No. 44 under the reference A17003.

## AUTHOR CONTRIBUTION

ASB, YL and RLG acquired funding; CJ, JLP, ASB, FM, and RLG designed the study; VC orchestrated the immunizations, longitudinal follow-up of the animals and biological sample collection and biobanking. CJ and JLP processed and biobanked biological samples; NDB generated plasma cytokine data; CJ and YF analyzed plasma cytokine data; YF stained samples for mass cytometry analysis; EML, and ASG acquired samples by mass cytometry; YF, JLP, NT, and ASB analyzed data; YF and JLP prepared the figures; YF and ASB wrote the original draft of the manuscript; YF, JLP, NT, RLG, and ASB reviewed and edited it. All the authors approved the submitted version.

## ACKNOWLEDGEMENTS

We deeply thank all the members of IMVA-HB/IDMIT, in particular Laetitia Bossevot, Benoit Delache, Nina Dhooge, Karine Gombert-Rannou, Orianne Lacroix, Sébastien Langlois, Julie Morin, and Jean-Marie Robert, members of the IDMIT core facilities ASW (Animal Welfare and Science), L2I (Immunology and Infectiology), and LFC (FlowCytech).

## CONTRIBUTION TO THE FIELD STATEMENT

We have previously reported, using an animal model highly relevant for human immunology, that a vaccine, when injected subcutaneously, triggered the presence of neutrophils better equipped to respond to a second vaccination long after the first one. Here we show that the intradermal injection of the same vaccine failed to induce such late neutrophil phenotypic modifications, although it mobilized neutrophils early after immunization. Single cells were analyzed by mass cytometry, a technique allowing their detailed characterization, and data from two vaccine studies were matched. Our contribution is twofold: (i) we demonstrate that the route of vaccine injection, likely through the modulation of the vaccine biodistribution and/or of the magnitude and quality of the early inflammatory/innate effector responses plays a role in the long-term innate immunological imprinting; (ii) we propose a simple method to compare existing and newly generated mass cytometry datasets post-clustering.

## SUPPLEMENTARY FIGURE LEGENDS

**Figure S1. Control samples**. The same fixed and frozen control samples were stained and acquired with the samples from the vaccinated animals after *ex vivo* restimulation with a mixture of TLR ligands. **(A)** Gating strategy to define the CD66^high^ HLA-DR^-^, CD66^-/mid^ HLA-DR^-^, and CD66^-/mid^ HLADR^+^ cell populations. The non-stimulated control sample for the first staining/acquisition session is shown. **(B)** Comparison of the staining profiles using overlaid histograms.

**Figure S2. SPADE tree annotation**. The topology of the SPADE tree is shown, with the color of each node of the SPADE tree representing the median expression of the indicated markers among all cells from the entire dataset with a scale adapted for each marker. This allowed the identification of the major blood cell populations, including granulocytes. The size of the node is not proportional to the number of cells it contains.

**Supplementary Table 1.**
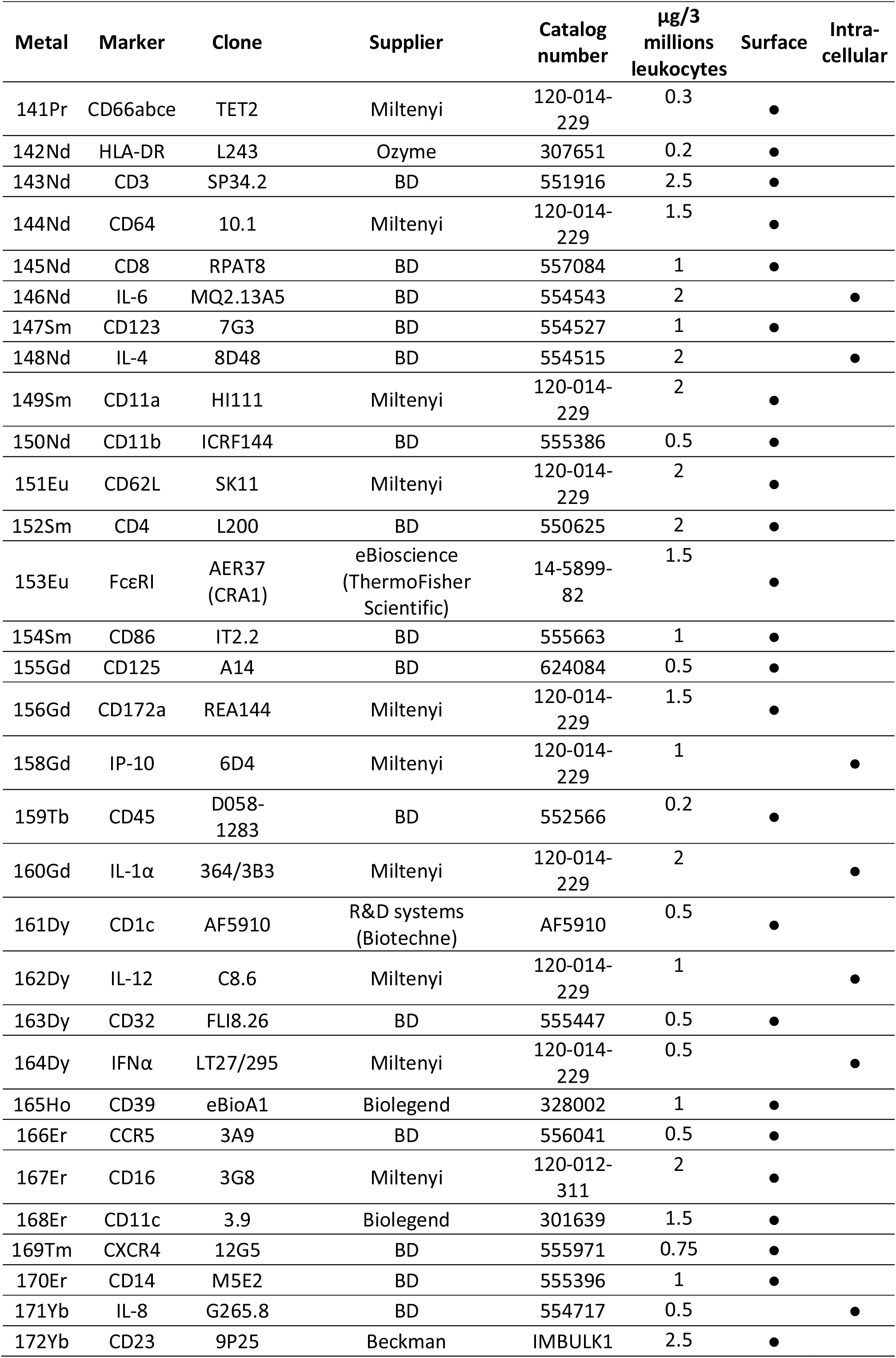
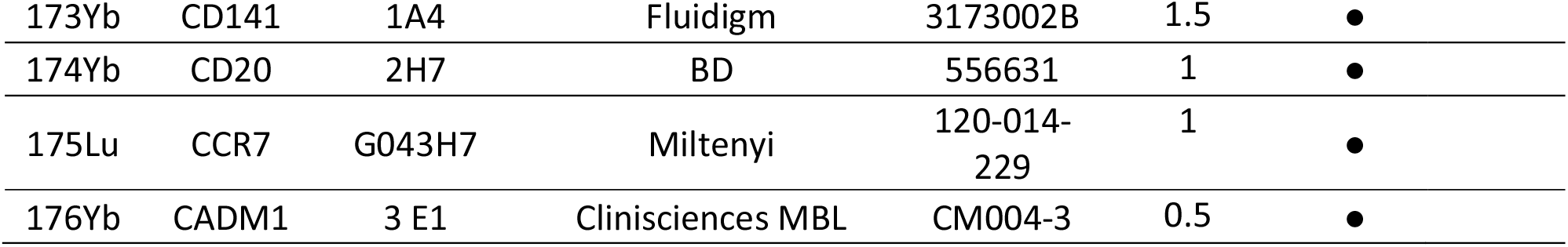
Antibody panel for mass cytometry.

