## Supplementary figures and images for "The route of vaccine administration determines whether blood neutrophils undergo long-term phenotypic modifications"

### Supplementary figures

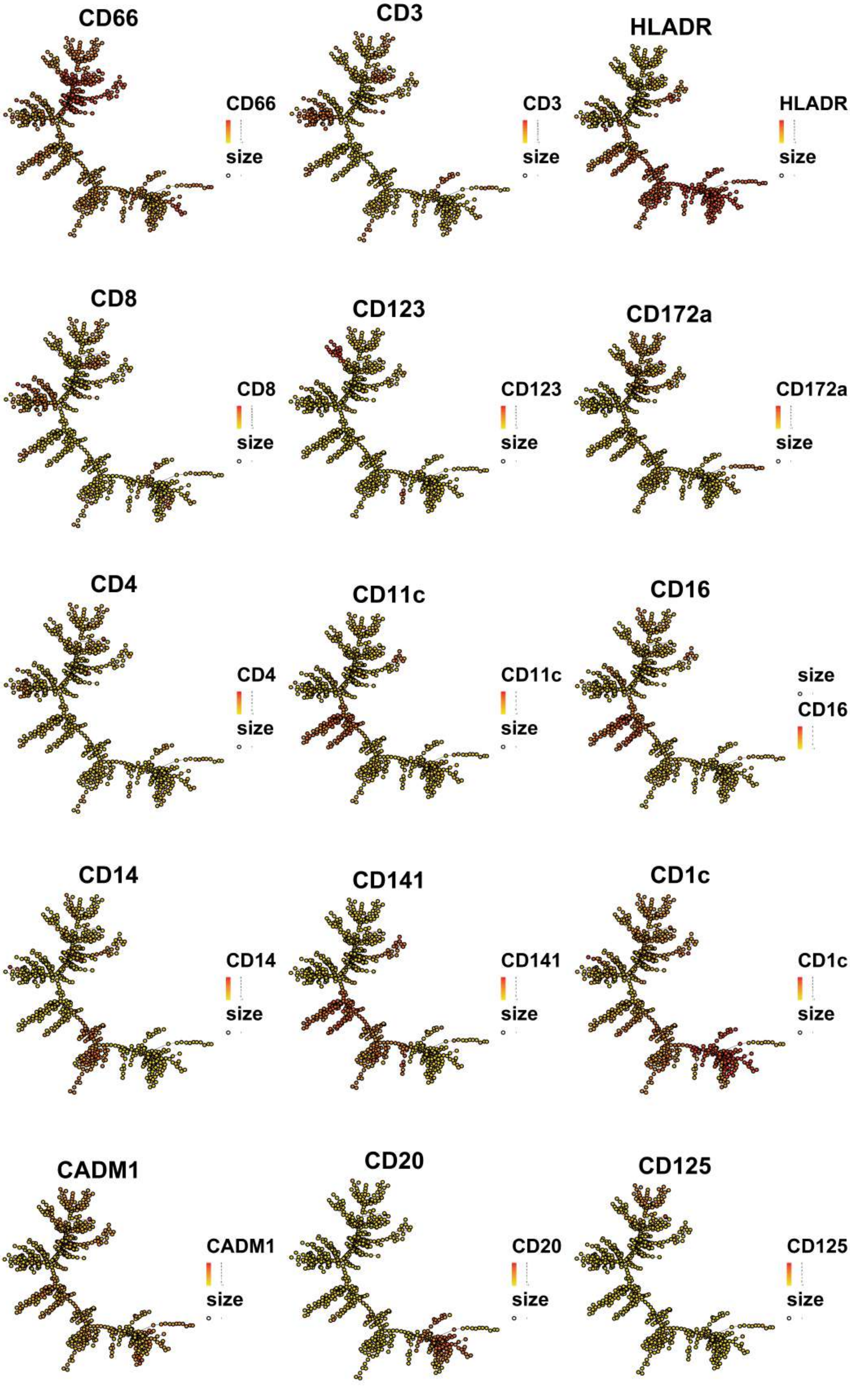

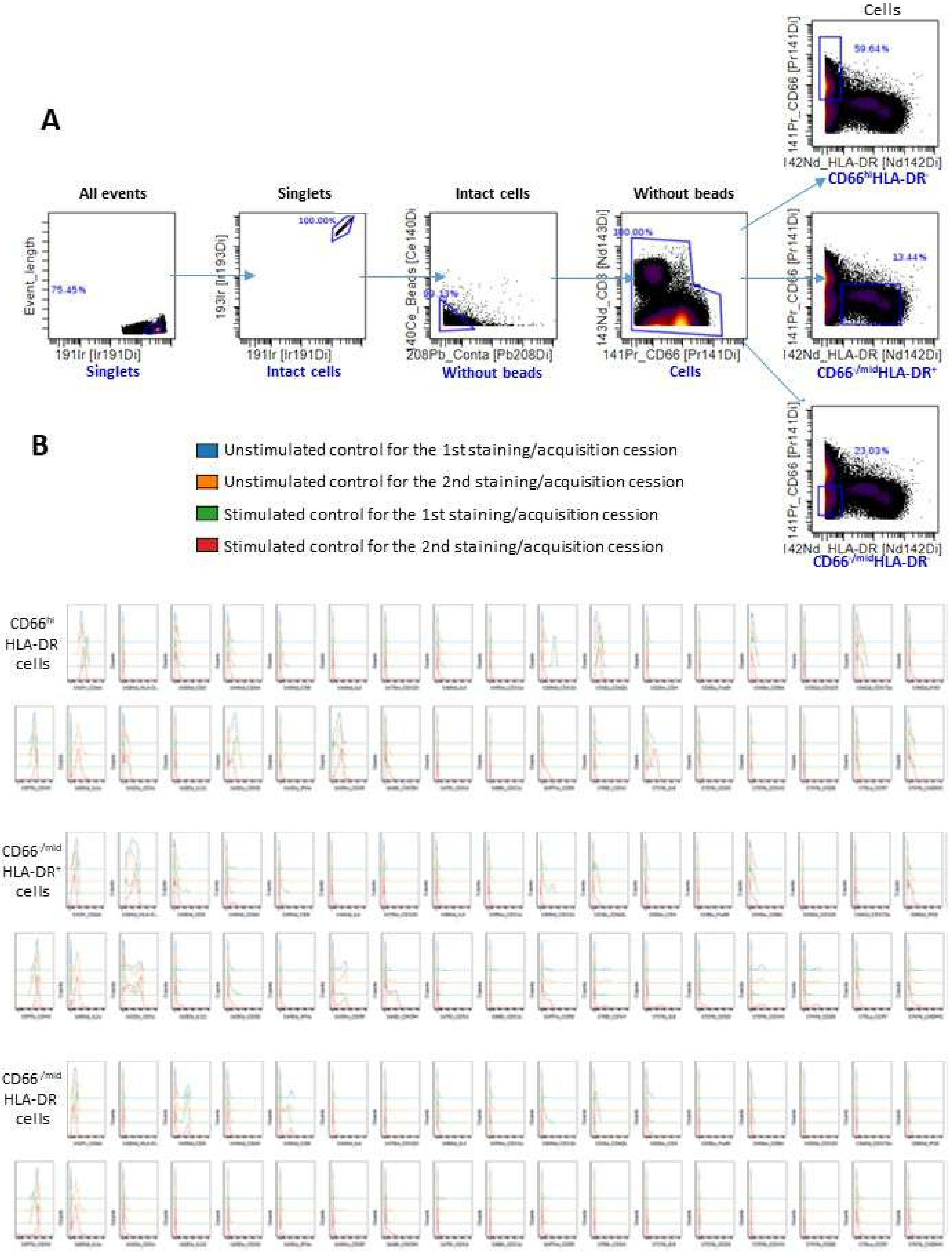
